# Probing single cell fermentation flux and intercellular exchange networks via pH-microenvironment sensing and inverse modeling

**DOI:** 10.1101/2022.05.03.490288

**Authors:** V. Onesto, S. Forciniti, F. Alemanno, K. Narayanankutty, A. Chandra, S. Prasad, A. Azzariti, G. Gigli, A. Barra, A. De Martino, D. De Martino, L.L. del Mercato

## Abstract

The homeostatic control of their environment is an essential task of living cells. It has been hypothesized that when microenvironmental pH inhomogeneities are induced by high cellular metabolic activity, diffusing protons act as signaling molecules, driving the establishment of cross-feeding networks sustained by the cell-to-cell shuttling of overflow products such as lactate. Despite their fundamental role, the extent and dynamics of such networks is largely unknown due to the lack of methods in single cell flux analysis. In this study we provide direct experimental characterization of such exchange networks. We devise a method to quantify single cell fermentation fluxes over time by integrating high-resolution pH microenvironment sensing via ratiometric nanofibers with constraint-based inverse modeling. We apply our method to cell cultures with mixed populations of cancer cells and fibroblasts. We find that the proton trafficking underlying bulk acidification is strongly heterogeneous, with maximal single cell fluxes exceeding typical values by up to 3 orders of magnitude. In addition, a crossover in time from a networked phase sustained by densely connected “hubs” (corresponding to cells with high activity) to a sparse phase dominated by isolated dipolar motifs (i.e. by pair-wise cell-to-cell exchanges) is uncovered, which parallels the time course of bulk acidification. Our method promises to shed light on issues ranging from the homeostatic function of proton exchange to the metabolic coupling of cells with different energetic demands, and paves the way for real-time non-invasive single cell metabolic flux analysis.

## Introduction

Fermentative processes are among the main modes of harnessing energy by cells. Despite their importance and the time elapsed since their discovery, they still continue to puzzle in regard to their basic function and mechanisms [1]. Since the first observation of fermentation inhibition by oxygen [2] and given their substantially lower efficiency with respect to oxidative pathways, fermentative pathways were initially seen as evolutionary relics with a subsidiary role, to be employed mainly in anoxygenic conditions and with problematic waste by-products. Subsequent observations, however, established their ubiquitous usage, even in presence of oxygen and especially for high energetic loads (e.g. during fast cellular growth), a phenomenon now known as “overflow metabolism” [3]. Because of its universality, current research efforts have been devoted to understand fermentation mechanism and function, e.g. in terms of volume constraints [4, 5] and/or temperature homeostasis [6]. A fundamental aspect of fermentative activity lies in its ecological dimension, specifically in its capability to alter the cellular microenvironment and most notably the pH level.

Cells dispose of a number of mechanisms to control their internal pH, which varies significantly across compartments (from ca. 8 in the mitochondrial matrix down to about 5 in secretory granules [7]). Cytoplasmic pH is especially impacted by the membrane potential (which tends to let positive ions in and negative ions out) and by the cell’s metabolic activity. Most notably, energy production by glycolysis, which occurs at high rates e.g. in cancer, generates cytosol-acidifying lactate. To counter this tendency to reduce cytosolic pH, cells utilize an array of transporters that couple proton export to the export (co-transporters) or import (exchangers) of some other metabolite, and whose activity is kinetically regulated by sites acting effectively as pH sensors [7]. This induces a reduction of the pH of the surrounding medium, which is in turn sensed by nearby cells. To maintain a stable extracellular proton level, then, these cells can respond by activating specific H^+^-sensitive channels and receptors.

The excretion of lactate by cells with large energetic demands and carbon consumptions, like tumors (so-called Warburg effect [8]) or neurons [9], constitutes an especially interesting case of microenvironmental acidification. It has indeed been hypothesised that lactate might become a fundamental vector of metabolic coupling within cellular populations in higher organisms [10, 11]: lactate-importing cells (acceptors) could rely on lactate-secreting cells (donors) for subsistence, while providing an essential bioremediation function by removing an acidifying metabolite from the microenvironment [12, 13]. This idea adds an ‘ecologic’ dimension to pH-driven intercellular communication [14].

While molecular mechanisms behind proton sensing, export and import are by now rather well characterized in a number of systems, due to their pathophysiological role for tumorigenesis and in the mammalian brain [15,16], much less is known about the actual intercellular proton-exchange network that is established. The major obstacle to overcome concerns the quantification of proton exchange fluxes for single cells within a population. Techniques to quantify cellular metabolic fluxes are in general well developed for the bulk of (mostly microbial) cell cultures [17–19]. Singlecell metabolomics, on the other hand, is less developed [20], with the exceptions of the growth rate [21], glucose uptake [22] and more in general nanoSIMS-based analyses [23,24], whose destructive character is however a serious shortcoming.

In this study, we gather information about the proton-exchange network by combining pH microenvironment sensing via recently devised pH-sensing ratiometric hybrid organic nanofibers [25–27] with constraint-based statistical inference. A sketch of the complete setup is shown in Figure 1. The first step involves the electrospinning onto glass slides (10 × 10 mm^2^), positioned on a custom rotating collecting system, of a 10% (w/w) polycaprolactone (PCL) chloroform/DMSO solution mixed with spherical and monodispersed SiO_2_ microparticle-based pH sensors (Figure 1a-b). In the second step, pancreatic ductal adenocarcinoma cells (PDAC, cell line AsPC-1) and pancreatic stellate cancer-associated fibroblasts (CAFs) are seeded onto the hybrid nanofibers and the fluorescence response of the pH-sensors during cell culture is recorded via time-lapse confocal laser scanning microscopy (CLSM) (Figure 1c). The third step involves precise automated quantification of intercellular proton (H^+^) exchange through physically constrained statistical inference (Figure 1d).

**Figure 1:**
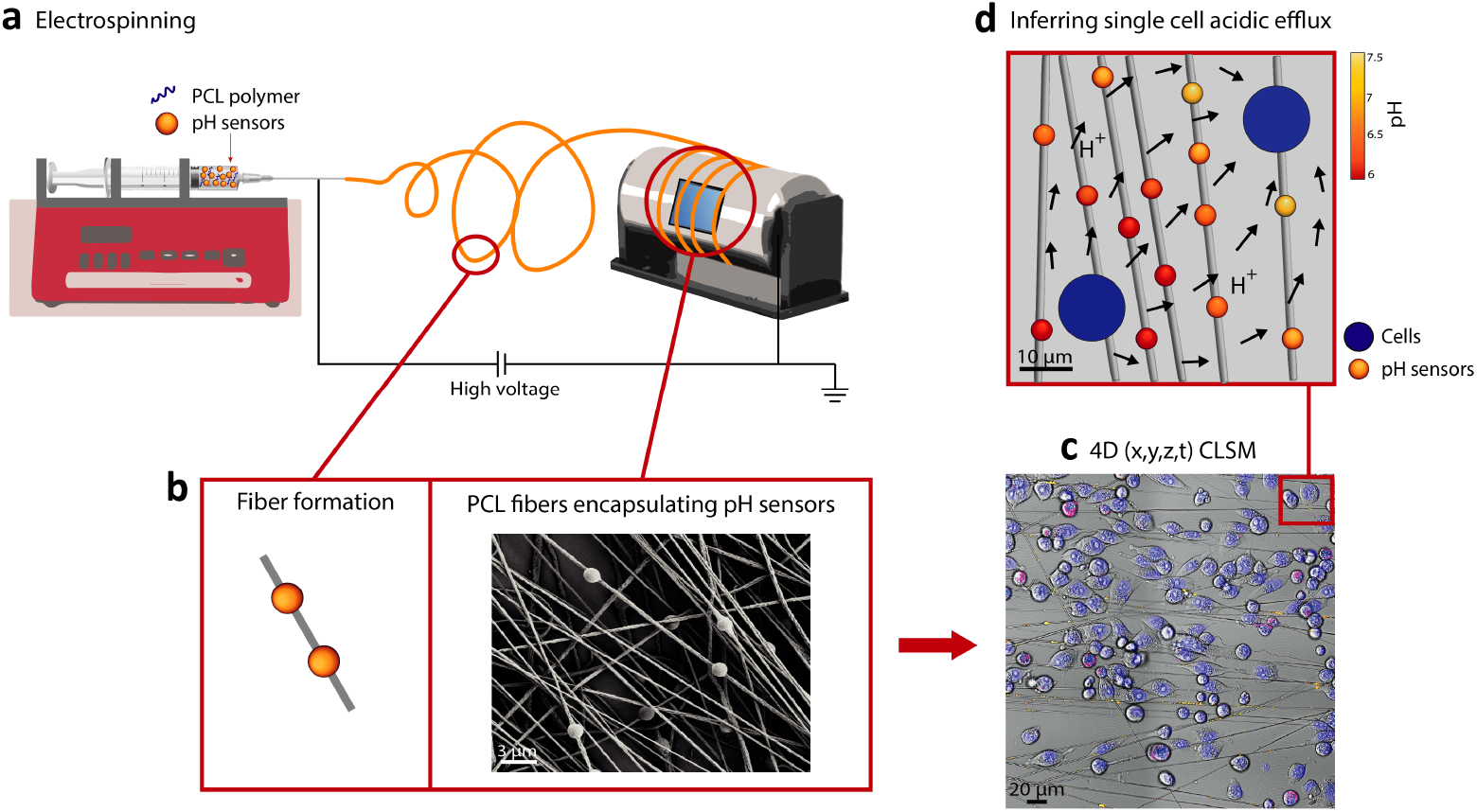
Schematic illustration of the workflow. (a) Sketch showing the fabrication of electrospun polycaprolactone (PCL) fibers embedding ratiometric SiO_2_-based microparticle sensors. (b) Representative SEM micrograph showing the morphology of PCL nonwoven mat of fibers carrying embedded pH sensors. (c) Representative CLSM image showing cells co-cultured on pH-sensing fibers and analyzed by CLSM time lapse imaging (*x*, *y*, *z*, *t*; *t* = 6h) (nuclei are shown in blue color, cell membranes are shown in magenta color for tumor cells). (d) Following spatial tracking of cells and probes, the whole pH gradient and the boundary single cell fluxes are reconstructed through physically constrained statistical inference.

Besides the accuracy of reconstructed fluxes, our method has the advantage of maintaining the sample intact. One can therefore monitor real-time single-cell behaviour potentially up to the whole population scale.

After illustrating our screening platform, we will focus on a concrete application, namely the reconstruction of the proton exchange network that underlies bulk acidification in mixed populations of CAFs and PDACs (Warburg effect). These results shed light on the nature of the ecology of cancer metabolism and, more in general, of metabolic overflow, while our method stands potentially as a first step towards non-invasive single cell metabolic flux analysis.

## Results

### Fabrication of pH-sensing scaffolds

We developed fluorescent pH-sensing nanofiber scaffolds composed of optical pH-sensors microparticles, highly suitable for monitoring pH changes in the surrounding environment [28–35], and polycaprolactone (PCL) nanofibers, because of their processability, biocompatibility, biodegradability, and mechanical stability [36–41].

The morphology of the hybrid pH-sensing fibers was studied in detail by means of scanning electron microscopy (SEM) and CLSM. The SEM images in Figures 2a,b show random nonwoven mat of PCL nanofibers bearing spherical pH-sensors aligned along the fiber longitudinal axis. The fibers present a typical rough and porous surface structure (Figure 2b), likely due to the use of chloroform/DMSO binary solvent system [42]. The PCL solution, as well as the concentration of the pH-sensors and the electrospinning conditions were adjusted in order to obtain uniform and bead-free nanofibers with a controlled diameter of 272.43 ±7.95 nm (Figure 2c). Notably, these nanosized electrospun fiber scaffolds provide a large surface-to-volume ratio, which is known to enhance key cellular functions, including adhesion, proliferation, and differentiation [43–45]. Moreover, their nanofibrous structure mimic the naive extracellular matrix (ECM), which plays a pivotal role in cell polarity as well as in cell-to-cell/matrix interaction [46–49]. Representative CLSM images of the pH-sensing nanofibers are shown in Figures 2d-f. Each pH-sensor microparticle is clearly detectable thanks to the FITC (Figure 2d) and RBITC (Figure 2e) dye molecules covalently linked with APTES to the surface of silica (SiO_2_) microparticles [50]. The FITC and RBITC fluorophores act as pH indicator and reference dyes, respectively, to enable ratiometric measurements of pH [51–54]. Thanks to the surrounding polymeric matrix, the pH-sensors remain stably immobilized into the lumen of the nanofibers during imaging, making pH-sensing ratiometric hybrid nanofibers an ideal biomaterial scaffold for monitoring local microenvironment proton changes in a fast and noninvasive way, with high spatial control and resolution. The pH-sensing nanofibers were used to culture pancreatic cancer cells (AsPC-1) and pancreatic stellate cancer associated fibroblasts (CAFs) and to monitor extracellular pH changes via time lapse CLSM acquisitions for 6h with time intervals of 10 minutes (Figure 2g). CLSM acquisitions were analyzed through segmentation algorithms in order to identify cells and sensors within the images (Figure 2h), to quantify FITC/RBITC fluorescence intensity ratio, hence the pH read-outs. Knowing the positions of the cells releasing or intaking acids and pH and positions, the acids efflux from each cell were inferred (Figure 2i), as described in detail in the following sections.

**Figure 2:**
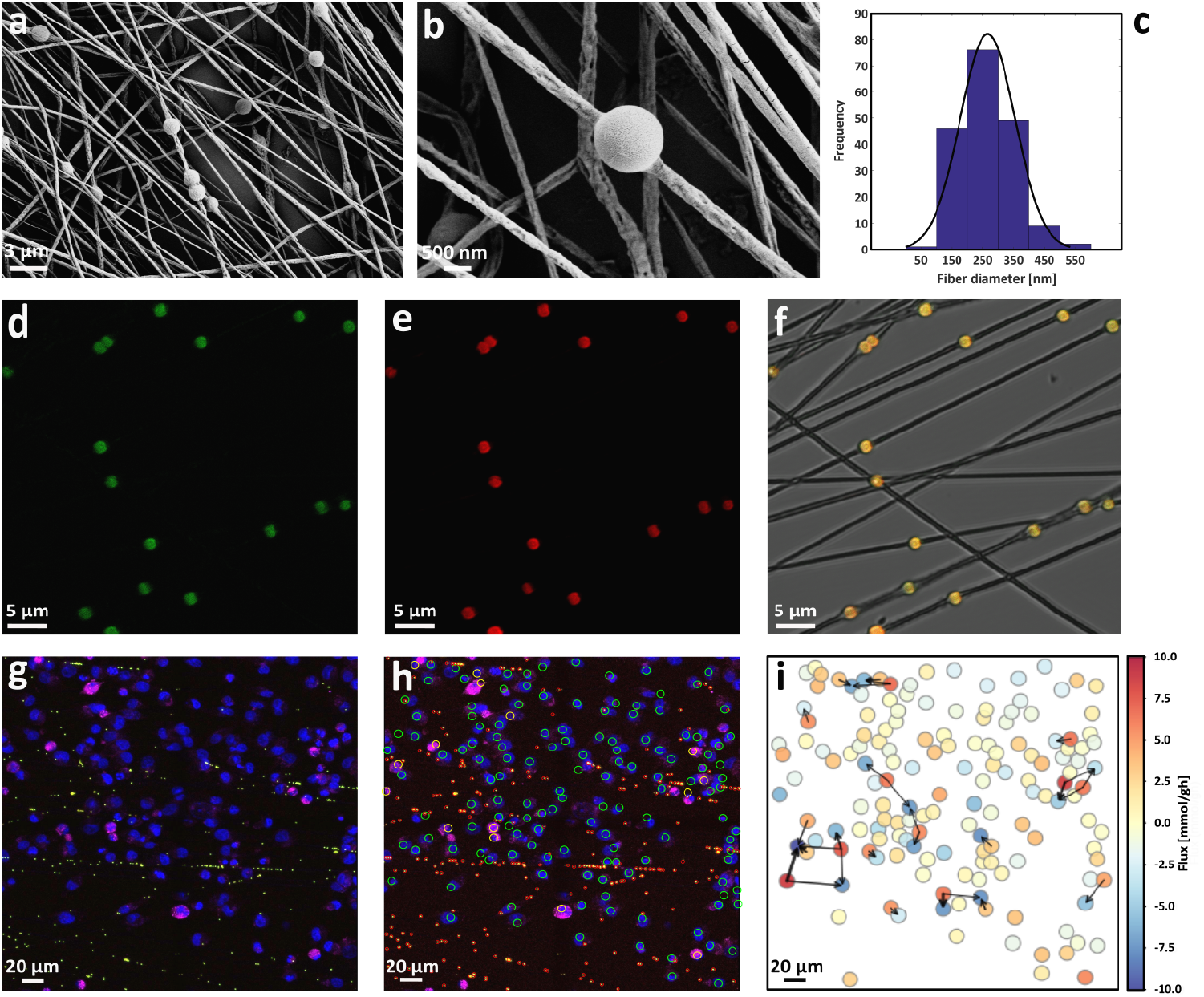
Scanning electron microscopy (SEM) micrographs showing (a) the pH sensors into the fiber’s lumen and (b) the corrugated morphology of the surface of individual fibers. (c) Graph illustration of the diameter distribution of the hybrid nanofibers. The superimposed continuous line is the best-fitting Gaussian curve. (d-f) Representative CLSM micrographs showing PCL nanofibers embedding pH sensors. FITC (green channel), RBITC (red channel) and overlay with bright field (BF, grey channel) are shown. (g) Representative images of CLSM time lapse image (maximum intensity projection) at the time point *t* = 3h, showing pH-sensing particles (FITC, green; RBITC, red), AsPC-1 cells (Hoechst, blue) and CAF cells (Hoechst, blue; Deep Red, magenta). (h) Result of the segmentation of the experiment in (e) showing the detection of the single pH sensors (red circles), AsPC-1 cells (green circles) and CAF cells (yellow circles). (i) Reconstruction of the cell fluxes through physically constrained statistical inference, with relative colormap.

### Inferring single-cell fluxes via pH landascape modeling

Measurements performed on the cell culture at any given time point yield (a) values of the pH (minus log-concentration of protons) at *M* locations and (b) the positions of *N* cells in a square region of linear size *L* = 500μm. Given (a) and (b), we want to determine the net proton exchange flux (import or export) for each cell. We solved this problem under a few simplifying assumptions. First, the proton concentration profile is taken to be stationary over our sample. This choice is motivated by the fact that experimental timescales (minutes) are much longer than the timescales over which concentrations are expected to equilibrate (seconds, assuming a diffusion coefficient *D* ≃ 7 × 10^3^ *μ*m^2^/s [55]) in regions of size L. In turn, stationarity implies that concentration profiles solve the Laplace equation ∇^2^*c*(**r**) = 0. Our second assumption is that the solution of the Laplace equation, i.e. the proton concentration at position **r**, can be written as

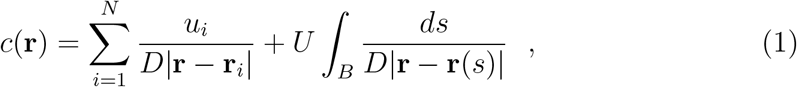

where **r**_*i*_ denotes the position of cell *i* (*i* = 1,…, *N*) and the parameters *u_i_* represent the net single cell fluxes we want to infer. The second term on the right-hand side models the flux *U* from the boundary *B* of the observed frame, whose value is to be inferred along with the *u_i_*s.

Equation (1) corresponds to the first terms in a multi-pole expansion of the solution of Laplace’s equation [56]. The truncation is justified as long as one is interested in the net exchange of protons with the medium by each cell. Higher-order terms in the expansion allow in principle for more refined descriptions (e.g. including separate import and export fluxes, shuttling of molecules inside cells, etc.). However, the larger number of parameters to be estimated would require a much more intensive sampling of the pH profile. By taking (1), we effectively focus on the metabolic generation and/or consumption of acids and on the ensuing symport of hydrogen ions.

We will now denote by *c_μ_* ≡ *c*(**r**_*μ*_) the proton concentration measured by the probe at position **r**_*μ*_ (*μ* = 1,…, *M*) and define the matrix of inverse distances 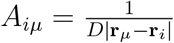. To simplify the notation, we include in the flux vector and in the distance matrix the terms due to the boundary of the frame, i.e. *u_0_* = U and 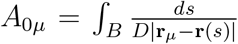. We can thus define for each frame *t* a cost function

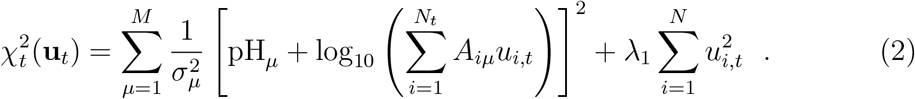

where, in the first term on the right-hand side, pH_μ_ = −log_10_c_*μ*_ denotes the pH level measured by the probe at position **r**_*μ*_, while *σ_μ_* represents the corresponding experimental error. The second term 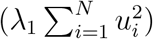, which enforces a Gaussian prior for the parameters u_*i*_, effectively imposes a uniform scale for the *u_i_s* (in agreement with the fact that metabolic fluxes are limited by physical constraints like intracellular and membrane crowding). This term is known as Tikhonov regularizer and it is necessary to prevent multicollinearity [57].

In order to avoid spurious effects in the inference due to variability in the number of cells in the frame during the experiment (where cells die, divide and migrate, both in and out of the observed frame), we finally consider a total cost function composed by the sum of the cost functions of each frame plus two additional cost terms, one imposing a limit to flux change for each cell across frames, and another constraining the overall mean flux 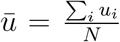 to the measured bulk value *u_b_* at each time point *t*:

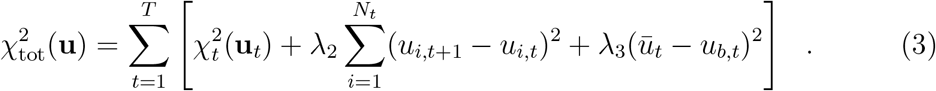

We will require that this weighted sum of residuals (the difference between empirical and reconstructed proton levels) is as small as possible. This is equivalent to assuming Gaussian-distributed residuals and inferred fluxes are maximum-likelihood estimates. Confidence intervals and errors on fluxes can be estimated accordingly from the inferred posterior. The minimization of 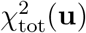 was carried out through an optimized Markov chain Montecarlo method (see methods and Supporting Information for details). Our inference scheme ultimately depends on the parameters λ_1_, λ_2_ and λ_3_, enforcing priors respectively for the intensity of single cell flux, its change in time and its average bulk value. Their estimate is described in the Supporting Information document.

Figure 3 displays a typical result from our experiments. Figure 3a,b,c shows the single cell fluxes (color scale in mmol/gdw/h ^1^); cells are schematically represented as disks of diameter 10 *μ*m in the same visual field at three different time-steps after cell deposition. Single cell fluxes appear to be evenly distributed in sign, i.e. roughly half of the cells secrete protons while the other half imports them. The quality of the reconstruction is illustrated in Figure 3d,e in space and time respectively. In Figure 3d the error between the reconstructed pH at the probes and the observed value is scattered against the observed value. More than 90% of the errors lies within the measured standard deviation (shades), although we detect a slight systematic deviation at low and high pH, that is however still within two standard deviations and amounts effectively to a smoothening of the gradient. In Figure 3e the comparison between reconstructed and measured pH in time is drawn for a given probe. In Figure 3f the bulk behavior of the pH (experimental, dots, and reconstructed, continuous lines) and the inferred bulk efflux (dashed line) is reported, while in Figure 3g the measured lactate levels in time that quantitatively correlate with the integral of the bulk efflux is presented. The bulk behavior is in good agreement with previous experimental findings [58] but single cell flux values are much higher. The bulk acidification observed during the Warburg effect therefore can be interpreted to be a mere leakage spilling from an intense network of cellular exchanges. In quantitative analogy with electrostatics, we highlight the presence of dipolar interaction motifs. The intensity of one motif as a function of time is shown in Figure 3h: the flux values of dipole-forming cells are mutually correlated and exceed the bulk value by three orders of magnitude (10mmol/gdw/h vs 10^−2^mmol/gdw/h). A general feature of the set of measured single-cell fermentation fluxes is its strong heterogeneity. This aspect has been analysed by building the empirical distribution as shown in Figure 3g. One sees that the tails strongly deviate from the superimposed Gaussian fit. This quantifies the intuition that a handful of cells carrying extreme fluxes are indeed responsible for a macroscopic fraction of the acidification level in the observed visual field.

**Figure 3:**
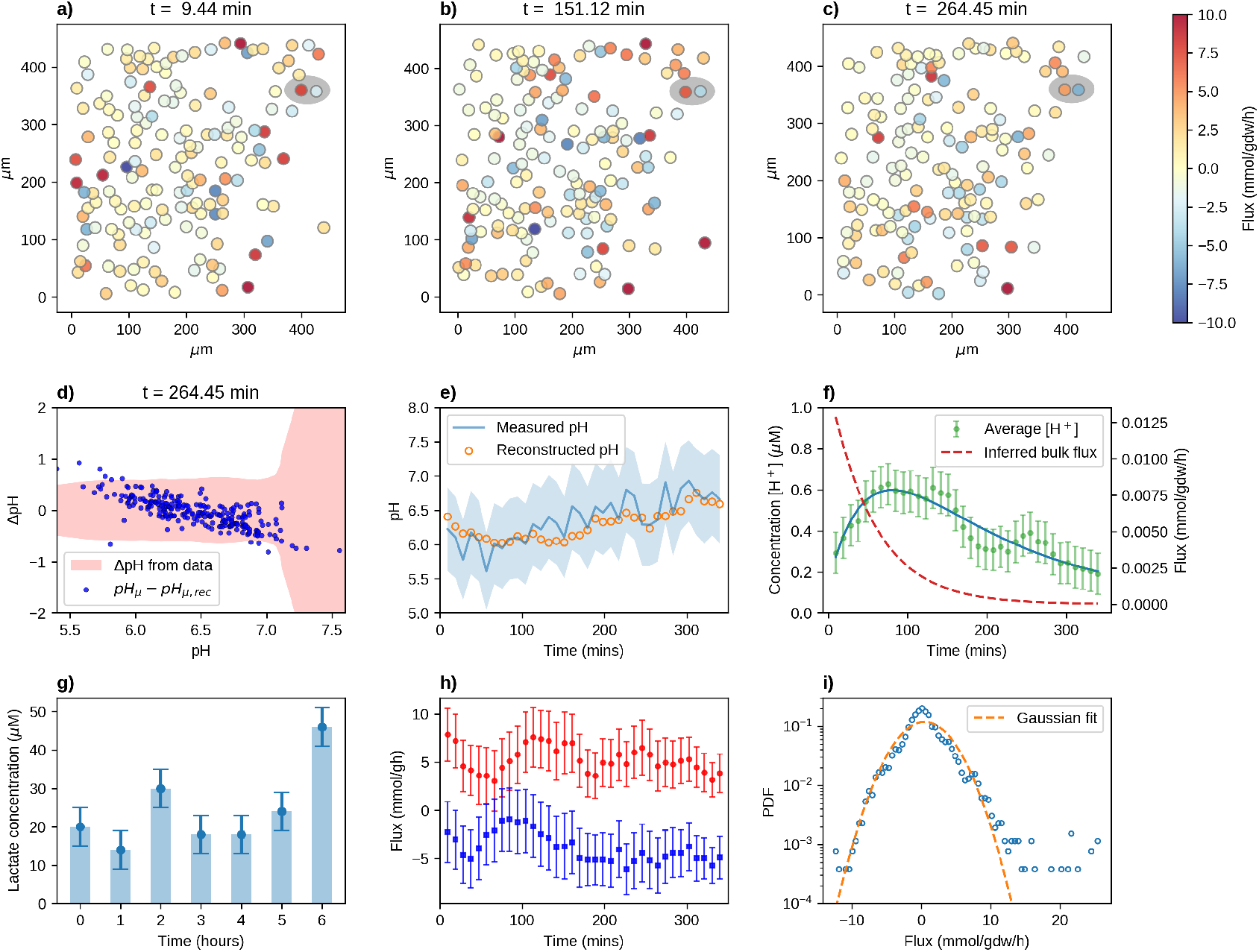
(a-c) Snapshots at different time points (at *t*_A_ = 9 min, *t_B_* = 151 min and *t_c_* = 274 min after the cell culture is settled) of the same square visual field (length *L* = 500 μm) during a typical experiment. Cells are represented schematically as disks of diameter 10 μm whose colour intensity scales with the flux (side bar, blue vs red for importing vs exporting flux respectively). Probes not shown. (d-e) quality of the reconstructed pH gradient profile. In (d) the error between the pH calculated from the inferred fluxes and the experimentally observed pH is plotted against the latter for each probe (at time *t_c_* = 274min). In (e) the time trace of the pH measured by a given probe is reported alongside the reconstructed trend in that spatial point. Shaded areas represent the experimental error on the pH at the probes. (f) Time trends of the bulk [*H*^+^] concentration (experimental, dots and reconstructed, continuous line, left *y* scale) and inferred bulk acidic efflux (dashed line, right *y* scale). (g) Time trend of the experimentally measured bulk lactate concentration in a biological replicate. (h) Single cell flux intensity (in mmol/gdw/h) as a function of time (in min, sampling every 10 min) of the cells forming the dipole motif highlighted in the upper right corner of the frames in (a-c). (i) Single-cell experimental flux distribution (in mmol/gdw/h, (dots) and its gaussian approximation (lines) in linear-logarithmic scale. The histogram is built from all single-cell flux values (100 – 200 cells per frame) and time-frames (36 frames resulting from a 6-hour experiment sampled every 10 minutes) tracked in one visual field of one experiment.

### Reconstruction of the cell-to-cell exchange network

In order to re-cast single-cell fluxes as pairwise exchange connections we observe that, in general, given a particle that starts to diffuse at **r**_0_ and *A* absorbing points at positions r*_j_* (*j* = 1,…, *A*), the probability that the particle is absorbed by one of them also satisfies the Laplace equation [59]. This means that, if *P*(**r**_*j*_|**r**_0_) denotes the probability that the particle initially at **r**_0_ is absorbed at **r**_*j*_, then

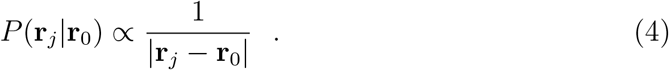

Based on this, we define the flux from cell *i* (with *u_i_* > 0) to cell *j* (with *u_j_* < 0) as

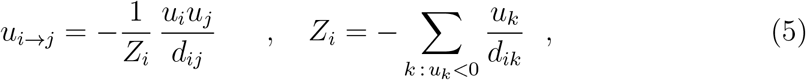

where *d_ij_* = |**r**_*j*_ – **r**_*i*_|. The term 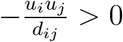 links the exchange between *i* and *j* to the magnitude of their proton fluxes and to how far i and j are from each other (the farther away they are, the less likely it is that they are connected as per (4)). The normalization factor *Z_i_* simply ensures that

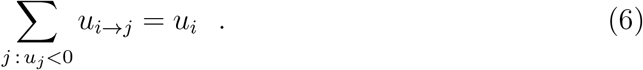

This defines a weighted directed network for our system that we can easily compute and analyse (see the Supporting Information for further details).

Figure 4a,b,c showcases the same three snapshot of the system displayed in Figure 3a,b,c, with the aforementioned network structure superimposed. Arrows are added between cells whose exchange exceeds our average sensitivity 0.5 mmol/gdw/h. The heterogeneity of fermentative phenotypes highlighted in Figure 3e,f is at the origin of the strong heterogeneities in the intensities of cross-feeding exchange fluxes, whose empirical distribution is reported in Figure 4d. One indeed sees that it spans three orders of magnitude.

**Figure 4:**
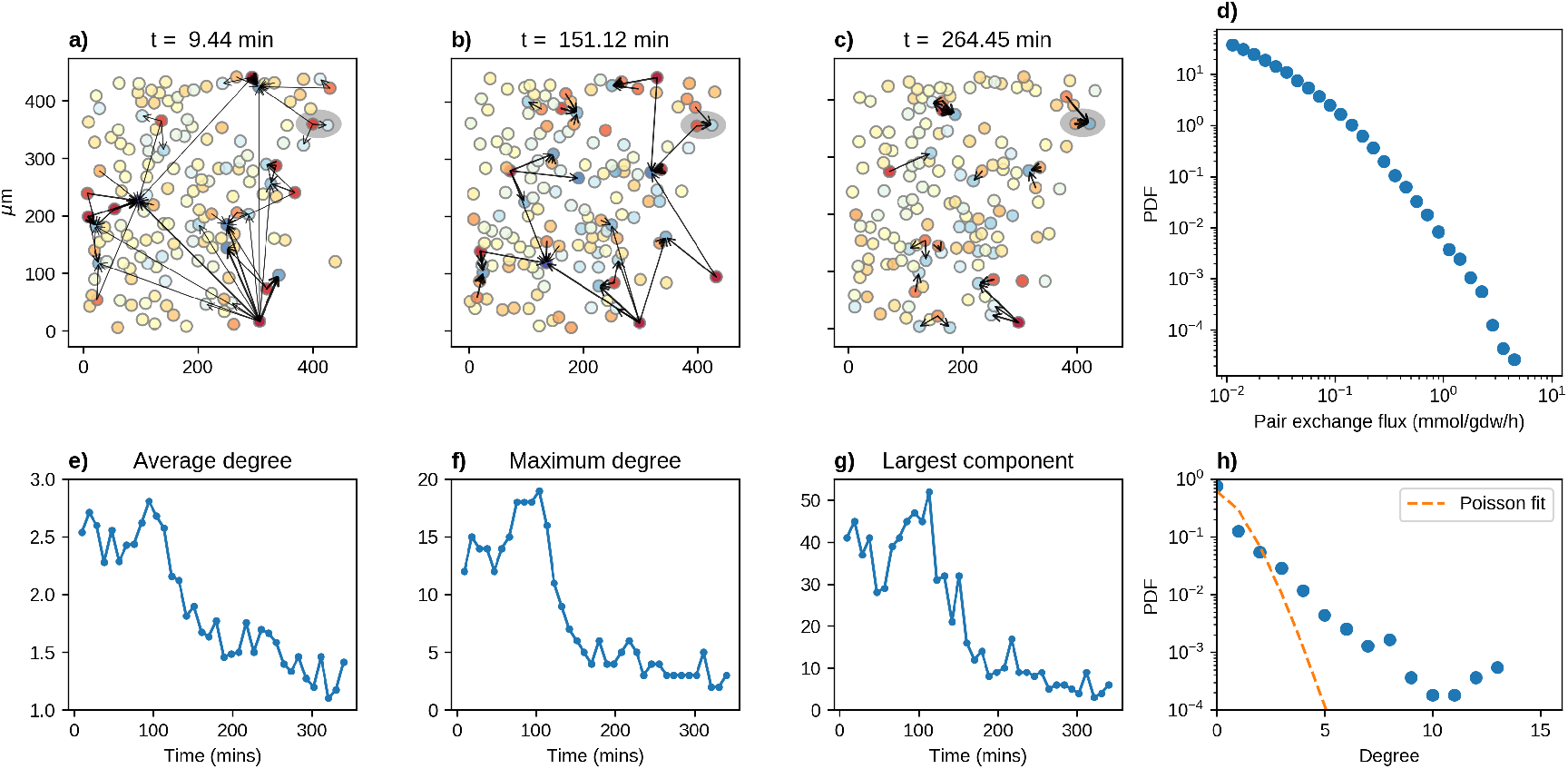
(a–c) Same snapshots of Figure 3a–c with superimposed inferred network structures. Arrows are drawn if the pairwise exchange flux exceeds 0.5 mmol/gdw/h, with a thickness proportional to flux intensity. (d) Distribution of pairwise exchange fluxes (in mmol/gdw/h) in double logarithmic scale (sampled over all frames). (e–g) Structural features quantifying the topology of the flux exchange network as a function of time (in min): average degree (e); degree of the node with maximum connectivity (f); and size of the largest connected component (g). (h) Degree distribution over all frames (dots) and corresponding Poissonian null hypothesis (same mean, lines).

We finally performed a standard graph theory analysis, in particular to calculate the time-dependence of the average degree 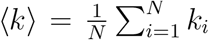, of the maximum degree, and the size of the largest connected component. These quantities are reported respectively in Figure 4e,f,g. Upon comparing Figures 4a and 4c, one sees that the trends highlight a crossover between qualitatively different regimes. At shorter times, a whole-frame-spanning exchange network is present, sustained by “hub” cells carrying high intensity fluxes. At longer times, such a network appears to dissolve, leading to a phase dominated by isolated dipoles. The topological structure develops in correlated fashion with the bulk pH acidification (Figure 3f). The presence of hubs can be appreciated upon looking to the experimental degree distribution depicted in Figure 4h as compared with a Poissonian null hypothesis of the same mean. The general trend is not significantly perturbed upon varying the threshold defining the links, that is the control parameter for the average network connectivity.

## Materials and Methods

### Chemicals

Polycaprolactone (PCL, molecular weight 80000 gmol-1, 440744, Sigma-Merck, KGaA, Darmstadt, Germany), chloroform (puriss. P.a., reag. ISO, reag. Ph.Eur., 99.0 – 99.4% (GC)-32211, Sigma-Merck, KGaA, Darmstadt, Germany), DMSO (dimethyl sulfoxide, BioUltra 99.5% GC-41639, Sigma-Merck, KGaA, Darmstadt, Germany) and ethanol (puriss. p.a., ACS reagent, Reag. Ph.Eur., 96% v/v-32294, Honeywell, USA) were used for the fabrication of electrospun fibers. Tetraethyl orthosilicate (TEOS, item code: 131903), (3-Aminopropyl)triethoxysilane (APTES, item code: 440140), potassium chloride (Item code: P9541), ammonium hydroxide solution 28% (item code: 211228), Fluorescein 5(6)–isothiocyanate (item code: 46950), Rhodamine B isothiocyanate (Item cod: R1755), tygon^®^ formula 2375 laboratory tubing I.D. x O.D. 1.6 mm x 3.2 mm (Item code: Z685585), were purchased from Sigma. 50 mL syringe was purchased from HSW HENKE-JECT^®^, NE-4000 syringe pump from New Era pump systems.

### Cell lines

Human pancreatic cancer cell line AsPC-1 (ATCC^®^ CRL-1682^™^) were obtained from American Type Culture Collection (ATCC, Rockville, Md., USA) and cultured at 37 °C in a humidified 5% CO_2_ incubator according to ATCC protocols. Cancer associated fibroblasts, CAFs (Vitro Biopharma Cat. n. CAF08) were cultured in DMEM medium (D5671, Sigma-Merck KGaA, Darmstadt, Germany) supplemented with 10% FBS (F7524, Gibco, Thermo Fisher Scientific), 2mM glutamine (G7513, Sigma-Merck KGaA, Darmstadt, Germany), 1% penicillin/streptomycin (P0781, Sigma-Merck KGaA, Darmstadt, Germany) and 1 μg/ml Recombinant Human Fibroblast growth Factor-basic (FGFb) (Catalog # 13256-029 Gibco, Thermo Fisher Scientific) at 37°C with 5% CO_2_. Cell lines were subcultivated with a ratio of 1: 5 and passaged 2 times per week. Mycoplasma contamination were routinely tested by Mycoplasma PCR detection kit (G238 abmGood, Canada).

### Image Analysis

Input data are raw images which, concretely, consist of information collected by three independent channels:

- Red Channel (particles emit a constant signal regardless of local pH)
- Green Channel (particles emit a signal proportional to local pH)
- Blue Channel (this is related solely to nuclei emission; hence it helps splitting cells from sensors)

Two algorithms were applied in series to all images as pre-processing steps, the former [Algorithm A] to identify cells or sensors within the image, the latter [Algorithm B] to quantify their intensity, hence their pH read-outs (for sensors only). More details are reported in the Supporting Information.

### Monte Carlo method

The inference setup described above is a maximum likelihood problem where fluxes are assumed to be distributed as 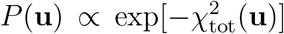 and the optimal guesses for **u** are those that maximize the probability of occurrence. Points of the posterior distribution have been sampled with an optimized Monte Carlo method based on the Metropolis-Hastings algorithm [60], i.e. by definining a Markov chain in the flux space based on a random walk whose steps have conditional probabilities 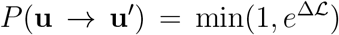, where 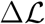 is the variation of log-likelihood 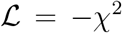 in going from **u** to **u**′. An analytical gaussian approximation of the log-likelihood rate function has been used to over-relax the random walk to tackle ill-conditioning and provide a warm start. The maximum likelihood optimal configuration has been found through simulated annealing while confidence intervals and errors have been estimated numerically by exploiting the invariance properties of the log-likelihood functions. More details are reported in the Supporting Information while codes implementing the method are available at https://github.com/demartid/infer_single_cell_fermentation_codes_data

## Conclusions

In this work we proposed and tested a method for the measurement of single cell fermentation fluxes, based on high spatial resolution cell seeded pH-sensor scaffolds and constraint-based statistical inference. The inverse character of the methodology makes it non-invasive and we applied it to follow in real time the acidification of the environment surrounding a cancerous population (Warburg effect) at the resolution of single cells, also in complex cellular systems, such as tumor and stromal cell cocultures. We highlighted the existence of a network of proton exchanges among cells in line with the lactate shuttle hypothesis. One of the most straightforward application of our method would be thus to probe the lactate shuttle hypothesis in physiological and pathological contexts, like the neuron-astrocyte and tumor-stroma metabolic partnership. The quantification of the exchange intensity reveals strong heterogeneity where a handful of cells is responsible for a large fraction of the pH gradient. Extreme single cell flux values are of the order of 10 mmol/gdw/h and overcome bulk values (roughly 10^-2^mmol/gdw/h) by at least 2 orders of magnitude. The former values are compatible with those measured in microbic overflows [61] while the latter are consistent with previous measurements of the (bulk) Warburg effect [58]. This would seem to suggest a common mechanism at work (e.g. saturation of physical constraints). In this regard it would be important to correlate the measurements of single cell fermentation with putative determinants of the overflow, like for instance single cell growth and/or oxidative rates. This would open the way for a non-invasive spatial metabolic flux analysis able to resolve the whole carbon flux at single cell resolution and the ensuing inter-cellular interactions. Such measurements will thus have significant impact on our understanding of the ecology and metabolism of cellular populations, in particular setting the experimental ground for recent quantitative theoretical approaches based on statistical mechanics [62–68].

## Acknowledgements

The authors gratefully acknowledge the ERC Starting Grant INTERCELLMED (project number 759959), the My First AIRC Grant (MFAG-2019, project number 22902), the Tecnopolo per la medicina di precisione (TecnoMed Puglia) Regione Puglia: DGR n.2117 del 21/11/2018, CUP:B84I8000540002. ADM acknowledges financial support from a Marie Skłodowska-Curie Action grant, GA No. 734439 (INFERNET). AB acknowledges financial support from Ministero degli Affari Esteri e della Collaborazione Internazionale, BULBUL grant (Italy-Israel). FA acknowledges financial support from Progetto PON R & I ARS01-00876 BIO-D “Sviluppo di Biomarcatori Diagnostici per la medicina di precisione e la terapia personaliz-zata”.

## Footnotes

1 The flux unit has been chosen to facilitate comparison with typical bulk values and assuming an average cell dry weight of 0.5ng

