## supplementary information for "Probing single cell fermentation flux and intercellular exchange networks via pH-microenvironment sensing and inverse modeling"

### 1 Further details on experimental methods

#### 1.1 Synthesis of particle-based pH-sensors

Ratiometric fluorescent pH-sensors based on silica microparticles were synthesized via recent work [1]. Briefly, dye molecules of FITC and RBITC were covalently linked with APTES in ethanolic solution for 3 hours in dark, 5 mg FITC and 6.5 mg RBITC were dissolved respectively in 3 mL anhydrous ethanol followed by addition of 13 $\mu$L APTES (20.6 mM). The two solutions of respective dyes with APTES were kept on magnetic stirrers for 4 hours at room temperature in dark. The formed FITC-APTES and RBITC-APTES conjugates were used directly in the next step without further purification. In the next step the silica microparticle buildup starts with the formation of seed suspension (Step 1) followed by its growth (Step 2) by slow addition of the monomer TEOS and dye-APTES conjugated molecules. The seed formation is achieved by dissolving 23 mg KCl in 9.45 mL of deionized water in a round bottom flask followed by addition of 96 mL of absolute ethanol, 6 mL of ammonium hydroxide (28%) and 1.73 mL of TEOS. The solution in the flask was continuously stirred using a magnetic bead at 240 rpm for 30 minutes. The next step involved increasing the size of the seed particles by slowly adding monomer TEOS along with dye conjugated APTES. Here 4.4 mL of TEOS, 2 mL of FITC-APTES and 2 mL of RBITC-APTES were dissolved in 33 mL of absolute ethanol. This solution was then slowly added drop by drop (flow rate: 0.05 mL/minute) into the seed solution that is being stirred at 240 rpm, using a 50 mL plastic syringe and tubing. During the seed formation and growth of the fluorescent silica particles the flask was kept airtight to using septum to prevent ammonia leakage. The reaction proceeded for 24 hours followed by careful pipetting of the supernatant, leaving larger debris of aggregated particles in the bottom. The collected particles in the supernatant were washed three times in ethanol using centrifugation at 2000 rpm followed by washing with deionized water thrice. The particles were stored in ethanolic suspension at 4°C.

#### 1.2 Fabrication and characterization of pH-sensing hybrid nanofibers

PCL solution for electrospinning was prepared using chloroform/DMSO in 90:10 v/v ratio. PCL was first dissolved in 900 $\mu$L of chloroform at a concentration of 10% (w/v), at room temperature with overnight stirring. 36 mg of pH-sensing microparticles were dispersed in 100 $\mu L$ of DMSO, transferred in an ultrasound bath (Digital Ultrasonic Cleaner, Argo Lab) at 20°C for 30 minutes to reduce aggregation, and finally added to 900 $\mu L$ of PCL solution in chloroform. After addition of the pH-sensing microparticles, the resulting solution (1 mL) was left stirring for 4 hours at room temperature. 1 mL of the solution was loaded in a 1 mL plastic syringe with 21G stainless steel needle and the syringe was placed on a syringe pump (E-fiber, SKE Advanced Therapies). A horizontal electrospinning set-up was used with a custom rotating collector having a rotating speed of 2200 rpm. The applied positive and negative voltages were +14 KV and −6 KV respectively, the tip of the needle to collector distance was 10 cm and the feeding rate was 0.8 mL/h. The hybrid organic nanofibers were collected on $1x1$ cm glass slides positioned on the target for 30 seconds. Before electrospinning, the glass slides were cleaned in a bath of ethanol for 5 minutes and dried with compressed nitrogen gas. All the experiments were conducted at room temperature ($19 - 22$°C) and a relative humidity of 35% within a closed chamber. Morphological characterization of the pH-sensing microparticles has been carried out by scanning electron microscopy (SEM, Sigma 300VP, Zeiss, Germany) analysis (accelerating voltage of 3 KV, secondary electron detector (SE2), 5000x, 10000x and 30000x magnifications). All the samples were sputter coated with an 8 nm thick gold layer (Safematic CCU-010 LV Vacuum Coating) prior to their observation under the microscope. SEM images were analyzed on ImageJ [2] software by drawing linear regions of interest at random locations to extract the average fiber diameter.

#### 1.3 Calibration of pH-sensing hybrid nanofibers

Calibration of pH-sensing matrices was performed by adding 200 $\mu$l of pH-adjusted cell medium (pH values $5.0, 6.0, 7.0, 8.0$) on the top of the nanofibers. Samples were allowed to equilibrate for 15 minutes, and then imaged using a confocal laser-scanning microscope (CLSM, SP8 Leica Microsystem, Mannheim, Germany) at $\lambda_{exc} = 488$ nm to collect the fluorescence signals in the green channel ($\lambda_{em} = 500 - 550$ nm) and at $\lambda_{exc} = 561$ nm to collect the fluorescence signals in the red channel ($\lambda_{em} = 570 - 620$ nm). At least four CLSM images (232.5 $\mu$m ×232.5 $\mu$m) were acquired for each pH point, with around 300 sensors in each image. Images were then processed as described in the following section. The equation of the fit calibration curve was used to estimate unknown pH experienced by the sensors in a region of interest.

#### 1.4 Cell co-cultures on pH-sensing hybrid nanofibers

Prior to the cell culture experiments, the pH-sensing nanofibers deposited onto 1x1 cm glass slides were sterilized by UV exposure for 30 minutes and placed into $\mu$-slide 4 well chamber slide (IBIDI GmbH, Grafelfing). AsPC-1 tumor cells and CAFs were counted by Trypan Blue dye exclusion and $4 \times 10^4$ cells were seeded in each well in the ratio 70% CAFs and 30% AsPC-1 tumor cells. In order to facilitate cell tracking by automated algorithms, both cell types were stained with a nuclear marker, Hoechst 33342 (B2261, Sigma Aldrich) 1:1000, and only tumor cells were stained with a plasma membrane marker, CellMask TM Deep Red (C10046, Invitrogen, ThermoFisher Scientific) 1:1000 for 20 minutes. After washing in PBS 1X, cell suspension was pipetted drop by drop on the pH-sensing matrices and then incubated over night at 37°C with 5% $CO_2$ to allow cell attachment.

#### 1.5 Extracellular lactate quantification

Changes in lactate concentration in cell culture medium were measured using the Lactate-Glo™ Assay Kit (Promega® Madison, WI, USA) according to the manufacturer's instructions. Briefly, AsPC1 tumor cells and CAFs were seeded on pH-sensing nanofibers as described in section 1.4. After cell attachment, the medium was changed in each well and left for 15 minutes to equilibrate. Each sample was collected every hour by diluting 5$\mu$L medium into 95$\mu$L of PBS. Cell culture medium collected from each experimental time point were then subjected to an enzymatic reaction coupling lactate oxidation and NADH production with a bioluminescent NADH detection system provided by the Lactate-GloTM Assay kit. Luminescence was recorded using a microplate reader (ClarioStarPlus, BMG Labtech). Medium without cells was used as negative control for determining assay background. The data generated were normalized by dividing the relative luminescence units (RLU) by the cell number obtained for each condition. Lactate concentration was calculated through a standard curve using a titration of lactate and all measured samples were within the linear range from 0 to 200$\mu$M. Data represent the average ($\pm$ SEM) of three independent experiments.

#### 1.6 Statistical significance

Data extracted from time-lapse experiments are the mean $\pm$ standard error of the mean (sem) of three independent experiments. Three SEM images were acquired for each magnification (5000x, 10000x and 30000x) from three different samples. Fiber diameters extracted from the SEM images are presented as mean $\pm$ sem of 100 individual measurements from each sample. Each time-lapse experiment was compared with the proper calibration curve to convert the fluorescence value to a precise pH value.

##### 1.7 Sensing of pH in the cell cultures over time

pH variations of cells cultured on pH-sensing nanofibers were measured over time for 6 hours with a time interval of 10 minutes. Before each time-lapse acquisition, calibration of the system composed of cells on pH-sensing matrices was performed by adding 200 $\mu$L of pH-adjusted Leibovitz L15 medium (cat. num 21083027, Gibco, Thermo fisher Scientific). Samples were allowed to equilibrate for 15 min, and then imaged by a CLSM Leica SP8 maintaining a temperature of 37 °C. 200 $\mu$L of L15 medium were added to the sample composed of cell co-cultures on pH-sensing matrices before starting pH monitoring. During time-lapse CLSM measurements, FITC was excited with $\lambda_{exc} = 488$ ($\lambda_{em} = 500 - 550$ nm), RBITC was excited with $\lambda_{exc} = 561$ nm ($\lambda_{em} = 570{-}620$ nm), Hoechst was excited with $\lambda_{exc} = 405$ nm ($\lambda_{em} = 415 - 500$ nm), Deep Red was excited with $\lambda_{exc} = 633$ nm ($\lambda_{em} = 640 - 750$ nm). The sections (size 232.5 $\mu$m $\times$232.5 $\mu$m) were collected with a $z$-step of 1.55 $\mu$m. Four CLSM images were acquired for each time point with a resolution of $696 \times 696$ pixels.

### 2 Further details on image analysis

#### 2.1 [A] Particle (cell or sensor) detection

Assuming particles are brighter than the background, at first we store the maximal intensity level of the image (that we call M), then we apply a spatial band-pass filter (BPF) with the band passing through the pixels [R/2, 3R/2] -R being the candidate radius of the particle and the bandwidth- obtaining a filtered image FI, then we further apply a dilatation [3] to the filtered image to obtain a second filtered image (DFI). Points accounting for centers of the particles then fulfill the conditions

$$(DFI = FI)and(DFI > M \cdot Tol) \tag{1}$$

where the parameter Tol (tolerance) ranges in $(0, 1)$: the first condition ensures that the pixel is a local maximum, the second condition guarantees that the luminosity of the candidate center is brighter enough for further processing [4].

#### 2.2 [B] Intensity Evaluation

To estimate the average luminosity I of each point, we calculate an integral over the radius of the particle using the formula

$$\bar{I} = exp[\frac{\sum_p I(p) log I(p)}{\sum_p I(p)}] \tag{2}$$

where I(p) is the intensity of the given pixel p. To track sensors procedure A is applied to the sum of the Green and Red channels in order to obtain the positions of all the particles within the image. Using as center of integration, time by time, all the candidate center positions previously detected, procedure B is then applied -separately to the red and green channels- obtaining, for each particle, the ratio R of their intensities R=I(red)/I(green): this ratio is then transformed, upon relying on the calibration curve, to a pH value. To track cells instead, the solely usage of procedure [A] suffices.

##### 2.3 Cell and probes tracking across frames

The recorded positions of individual cells and probes vary across time not only due to the motion of cells but also due to the lateral shifts during the imaging process. For tracking objects across time, we employed a simple method based on comparing the positions of individual objects across consecutive time points and finding the closest match: Particle $i$ of a time point and particle $j$ of another time point are considered to be the same particle if the separation $d_{ij}$ between them is a minimum w.r.t. both variation of $i$ as well as that of $j$. This method was considered sufficient since the maximum displacement of the objects, as observed from the images, across any one interval in time was smaller than the minimum separation between the objects in any single frame. There were instances of cells at the boundary leaving and re-entering the frame of imaging, and also cells crossing over each other during motion, at which point our algorithm fails to track these cells anymore. In case of probes, the tracking fails when the lateral shifts during imaging are comparable to the minimum separation between individual probes, and also when cells pass over any probes affecting the imaging of those probes. With this rather conservative algorithm, we were able to successfully track more than 65% of the cells and more than 80% of the probes.

#### 3 Further details on computational methods

##### 3.1 Gaussian approximation

We will describe here an analytically tractable approximation of the inference task. This will play a fundamental role for the Monte Carlo Markov chain to solve the full non-linear problem, in order to

1. Provide a warm start: the approximate maximum likelihood point will be taken as initial point for the sampling thus reducing the transient equilibration time.
2. Tackle ill-conditionining: the approximate covariance matrix will be used to define a coordinate transformation (or in practice perferential sampling direction) that round the space and reduce sampling times by the condition number, that is the ratio between the maximum and minimum eigenvalue.

For sake of simplicity in notation we will consider here the expression for one time frame, that can be easily generalized to the full time evolution. Let us consider the full non-linear likelihood function (for sake of simplicity without priors)

$$\chi^2_{NL}(\mathbf{u}) = \sum_{\mu} \frac{(pH_\mu + \log_{10}(\sum_i A_{i\mu} u_i))^2}{2\sigma^2_\mu} \tag{3}$$

Upon considering $c_\mu = 10^{-pH_\mu}$ we can approximate the residuals

$$pH_\mu + \log_{10}(\sum_i A_{i\mu} u_i) = \log_{10}(1 + \frac{\sum_i A_{i\mu} u_i - c_\mu}{c_\mu}) \sim \frac{\sum_i A_{i\mu} u_i - c_\mu}{c_\mu} \tag{4}$$

that shall be valid for small deviations from the expected value and leads to a Gaussian expression of the likelihood.

$$\chi^2_L(\mathbf{u}) = \sum_\mu \frac{(c_\mu - \sum_i A_{\mu i} u_i)^2}{2c_\mu^2 \sigma_\mu^2} \tag{5}$$

Notice the weights in agreement with standard error propagation formulae. Standard linear algebra manipulations lead finally to the optimal solution

$$u_j^* = \sum_i (B_{ij})^{-1} b_i \tag{6}$$

$$\text{where} \quad b_i = \sum_\mu \frac{A_{\mu i}}{c_\mu \sigma_\mu^2} \tag{7}$$

$$\text{and} \quad B_{ij} = \sum_\mu \frac{A_{\mu i} A_{\mu j}}{c_\mu^2 \sigma_\mu^2}. \tag{8}$$

Upon regarding the $\chi^2$ as the large deviation function for the posterior it is straighforward to obtain an analytical approximate expression for the covariance matrix of fluxes (we indicate by $\langle \dots \rangle$ the average)

$$\langle u_i u_j \rangle = (\mathbf{B}^{-1})_{ij} \tag{9}$$

This matrix could be used for errors estimation, but it highly overestimates them. In practice, its principal component analysis gives quantitative insights on the typical scales and it has been used to remove the ill-conditioning in the inference via preferential sampling direction in the Monte Carlo Markov chain.

###### 3.1.1 Enforcing positivity constraints of the reconstructed concentration profile

The gaussian approximation has the drawback that the reconstructed profile is not anymore positive definite (a condition that is verified by construction in the full non-linear case). This can be ameliorated upon enforcing ad-hoc constraints that defines a quadratic convex optimization problem. We can thus impose that the reconstructed values of the concentration at the location of the probes $c_{\mu,\text{rec}} = \sum_i A_{\mu i} u_i$ shall strictly be non-negative. This leads to the problem

$$\begin{aligned} \text{minimise} \quad & \chi^2_L(\mathbf{u}) \\ \text{such that} \quad & \sum_i A_{\mu i} u_i \geq 0 \end{aligned}$$

This is in the standard form of a convex quadratic program, the we solved using the Goldfarb-Idnani algorithm [5].

#### 3.2 Priors and lagrange multipliers

##### 3.2.1 Tikhonov regularizer for the scale $\lambda_1$

We considered a regularizer $\lambda_1$ setting the single cell flux scale. This is done to prevent multicollinearity and it is equivalent to assume a prior taking into account previous experimental knowledge (e.g. on enzyme kinetics). The introduction of regularizers as quadratic terms in the log-likelihood rate function does not change the form and character of the maximum-likelihood solution as it can bee seen in the following. Consider the simplified form

$$\chi^2_{\lambda_1}(\mathbf{u}) = \sum_{\mu}(c_\mu - \sum_i A_{\mu i}u_i)^2 + \lambda_1 \sum_i u_i^2 \tag{10}$$

We have

$$\begin{aligned}\chi^2_\lambda(\mathbf{u}) &= \chi^2(\mathbf{u}) + \lambda\mathbf{u}^2 \\ &= \mathbf{c}'^{\mathbf{T}}\mathbf{c} - 2(\mathbf{c}'^{\mathbf{T}}\mathbf{A})\mathbf{u} + \mathbf{u}^{\mathbf{T}}(\mathbf{A}^{\mathbf{T}}\mathbf{A} + \lambda\mathbf{I})\mathbf{u}\end{aligned} \tag{11}$$

which on comparison with the previous solution it gives the minima $\mathbf{u}^*$ as

$$\mathbf{u}^* = (\mathbf{A}^{\mathbf{T}}\mathbf{A} + \lambda\mathbf{I})^{-1}\mathbf{A}^{\mathbf{T}}\mathbf{c}' \tag{12}$$

$$= (\mathbf{B} + \lambda\mathbf{I})^{-1}\mathbf{b} \tag{13}$$

We have fixed $\lambda_1 = 2.5 \cdot 10^{-2}$ (mmol/gdw/h)$^{-2}$, in turn corresponding to a typical flux scale of 20mmol/gdw/h consistently with known values of the maximum rate $V_{max}$ of monocarboxylate transporters [6].

##### 3.2.2 Time continuity across frames $\lambda_2$

As we remarked above the number of cells in the observed visual field varies due to (mainly) migration, proliferation and death, with cells entering and exiting the field view. This can introduce spurious terms in the inference: for instance the flux of cells nearby the boundary in one frame that exit the visual field in the next frame could be artificially rea-assigned to nearby cells. In order to minimize the effect of such artefacts we imposed time continuity across frame for tracked cells (see above) with a term connecting the log-likelihood rate function of different frames

$$\lambda_2 \sum_{i=1}^{N_t}(u_{i,t+1} - u_{i,t})^2 \tag{14}$$

The value of $\lambda_2$ has been set to match the observed time gradient for the probes, that is within 10% of the maximum observed variation or around 1mmol/gDW/h per time frame (10m), in turn giving $\lambda_2 = 0.5$(mmol/gDW/h)$^{-2}$.

##### 3.2.3 Matching the bulk trend $\lambda_3$

Upon averaging over the probes we have a measurement of the bulk pH $\bar{c}(t)$ as a function of time. The dynamics generating the latter can be essentially split into two basic contributions: i) the average or bulk cell efflux $\bar{u}$ ii) the medium buffering capacity. Upon inferring the former, this can be used to further constraint our single cell flux measurements upon adding a term

$$\lambda_3(\frac{1}{N}\sum_i u_i - \bar{u})^2 \tag{15}$$

in the log-likelihood rate function. We have thus considered the following model for the dynamics in terms of linear perturbation around a steady state (we drop the bars for simplicity of notation), with parameters $k$, $c_e$ describing the medium buffering and $k_u$, $u_s$

$$\dot{c} = u - k(c - c_e) \tag{16}$$

$$\dot{u} = -k_u u \tag{17}$$

This linear system has a straighforward analytical solution

$$u = u_0 e^{-k_u t} \tag{18}$$

$$c = (c_0 - c_e) + c_e(1 - e^{-kt}) + \frac{u_0}{k_u - k}(e^{-kt} - e^{-k_u t}) \tag{19}$$

and its four parameters has been inferred by a simple local grid search. We obtain $k = 0.42 \pm 0.3\mathrm{h}^{-1}$ $k_u = 1.0 \pm 0.2\mathrm{h}^{-1}$ $u_0 = 0.015 \pm 0.003\mathrm{mmol/gdw/h}$. These values can be used to make a estimate for the lactate concentration ($\dot{c}_{lac} \sim u$)

$$c_{lac} \sim u_0/k_u \sim 30\mu M \tag{20}$$

that is in quantitative agreement with the experimentally observed ($\sim 20 - 70\mu$M). The inferred time trend for $u(t)$ has been then used to constrain the average flux in our inference and standard statistical considerations suggest that the lagrange multiplier shall be be set to $\lambda_3 = N\lambda_1 \sim 4$ (mmol/gdw/h)$^{-2}$, where $N \sim 150$ is the average number of cells per frame.

##### 3.3 The Monte Carlo Markov chain

The posterior probability distribution for the fluxes is

$$P(\mathbf{u}) \propto e^{-\chi^2_{NL}(\mathbf{u})} \tag{21}$$

In order to sample from this distribution we employed the well known Metropolis-Hastings algorithm: starting from an initial configuration $\mathbf{u}_0$, define the series $\mathbf{u}_l$

1. Upon perturbing $\mathbf{u}_l$ propose a new vector $\mathbf{u}_{l+1,p}$, calculate the likelihood variation $\Delta\mathcal{L} = \chi^2_{NL}(\mathbf{u}_{l+1,p}) - \chi^2_{NL}(\mathbf{u}_l)$.

2. Accept the new vector $\mathbf{u}_{l+1} = \mathbf{u}_{l+1,p}$ with probability

$$\min(1, e^{\Delta\mathcal{L}}) \tag{22}$$

otherwise keep the old one $\mathbf{u}_{l+1} = \mathbf{u}_l$

The efficiency of such method relies on properly defined proposal steps (e.g. avoiding ill conditioning) and on initial points that are as close as possible to the typical equilibrium ones (warm start). The proposal step has been chosen by performing a random walk over the eigenevectors of the covariance flux matrix in the gaussian approximation and choosing as starting point the maximum likelihood solution of the approximated gaussian rate function, i.e.

- The initial point is $\mathbf{u}_0 = \text{argmax} \quad \chi^2_L(\mathbf{u})$
- The proposed point is $\mathbf{u}_{l+1,p} = \mathbf{u}_l + \delta_k \mathbf{C}_k$, where $\mathbf{C}_k$ is the k-th eigenvector and $\delta_k$ is a gaussian random variable of zero mean and standard deviation equal to the k-th eigenvalue upon diagonalizing the inverse of the flux covariance matrix in gaussian approximation. The number k is selected uniformly at random.

This reduces sampling times by a factor approximatively equal to the conditioning number, i.e. the ratio between the largest and smallest eigenvalue of the covariance matrix, that in our case is around $10^4$, parelling similar analysis for the bulk metabolic networks [7]. The maximum likelihood estimate has been retrieved finally via simulated annealing [8], i.e. multiplying the log-likelihood by a fictitious (inverse) temperature $\beta$ in the Metropolis scheme, that changes the acceptance probability to

$$\min(1, e^{\beta \Delta \mathcal{L}}) \tag{23}$$

and then performing a Montecarlo dynamics with $\beta$ gradually increasing to high values.

##### 3.3.1 Errors and confidence interval

Once the maximum likelihood estimate for the fluxes has been calculated, their relative errors can be read off from the expression of the log-likelihood itself upon considering its meaning of for the posterior probability through Bayes theorem [9]. Confidence intervals $(\delta_-, \delta_+)$ for the fluxes corresponding to $\sim$ 97.6% probability ($2\sigma$ for the Gaussian) have been estimated by looking at flux variations that change accordingly the log-likelihood

$$\mathcal{L}_{max} - \mathcal{L}(\mathbf{u}_{-i}, u_{i,opt} \pm \delta_{\pm,i}) = -2 \tag{24}$$

while for the error bars we use $\Delta\mathcal{L} = -0.5$ and symmetrize. This follows from standard theory on non-parabolic error and confidence interval estimates [10].

The error on pairwise flux exchange can be propagated from single cell fluxes through the formula and assuming for simplicity independent errors for the fluxes we have the formula for the relative errors ( $\epsilon_i = \delta_i/|u_i|$ )

$$\epsilon_{i\to j} = \epsilon_i + \epsilon_j + \sum_{k:u_k<0} \epsilon_k \phi_{ik} \tag{25}$$

$$\phi_{ik} = \frac{u_k/d_{ik}}{\sum_{l:u_l<0} u_l/d_{il}} \tag{26}$$

### 4 Supporting files description

Files can be found at
`https://github.com/demartid/infer_single_cell_fermentation_codes_data`

#### 4.1 Data

We provide two data files

- `probes.dat` reports the position and measured pH value, with error for each probe at given timepoints.
- `cells.dat` reports the position and measured flux, with error and confidence intervals, for each cell at given timepoints.

#### 4.2 Codes

We provide two codes

- `gauss.py` takes in input the position and pH value (with errors) of probes and the position of cells for a given frame and it gives in output the maximum likelihood estimate of the cell fluxes and their covariance matrix in gaussian approximation.
- `sim_ann.cpp` accepts in input the inverse distance matrices of cells and probes for all frames, the gaussian estimated flux and covariance, the cell tracking and it gives in ouput the maximum likelihood estimate of the cell fluxes with errors and confidence interval.

### 5 Supplementary Figures

#### 5.1 Snapshot of inferred fluxes per single frames

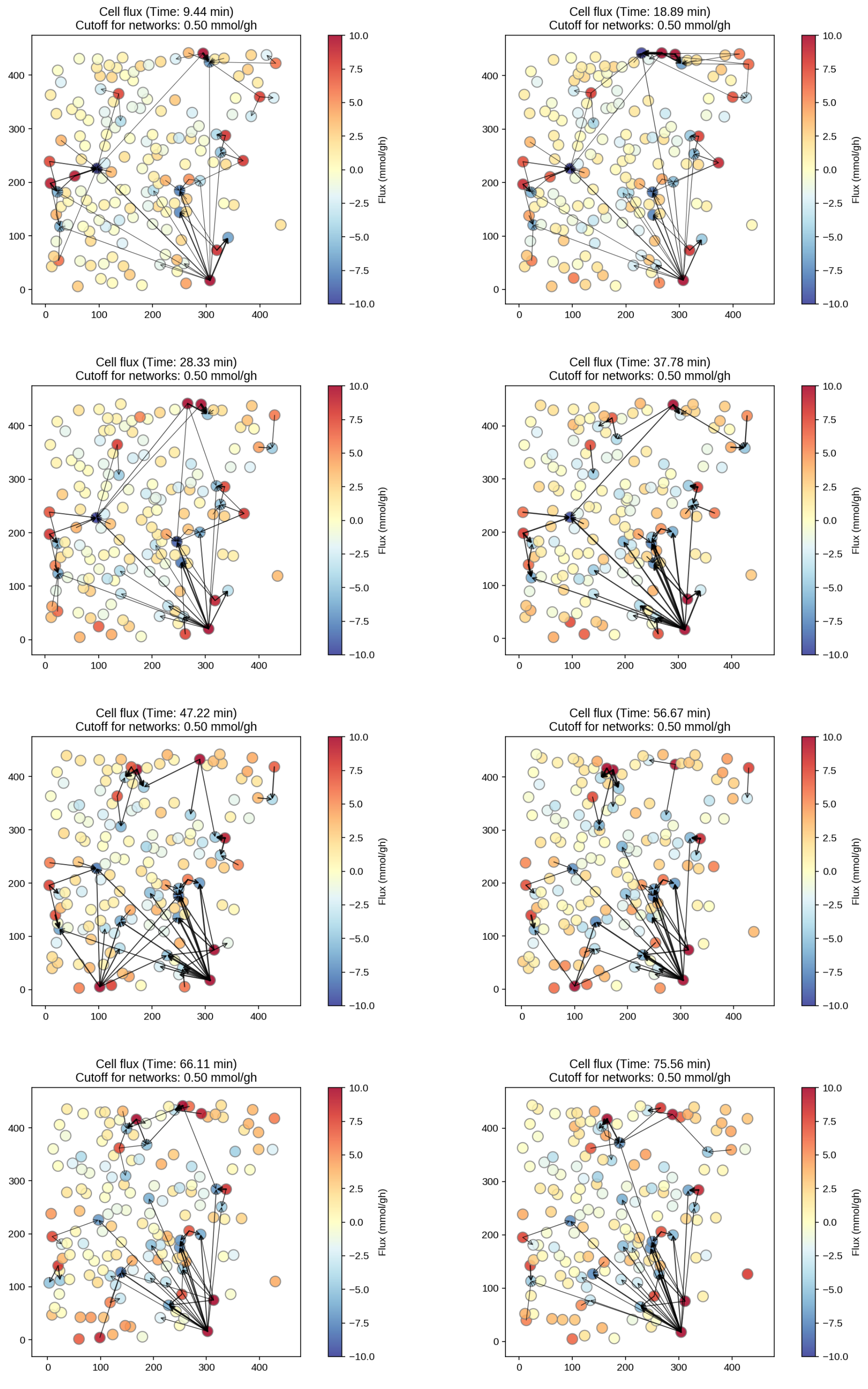

**Figure 1:** Cell fluxes time $t < 1h20m$.

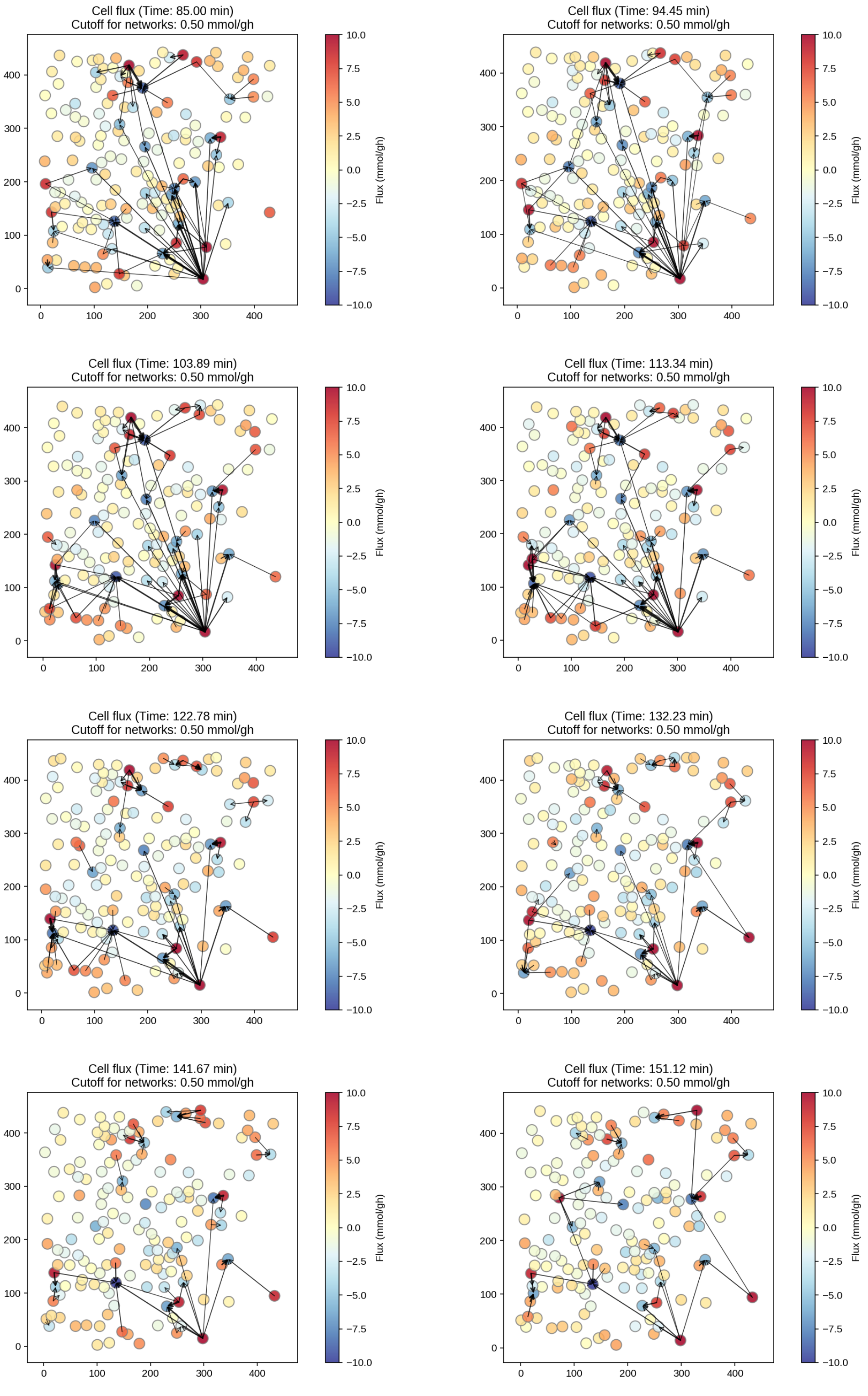

**Figure 2:** Cell fluxes time $1h20m < t < 2h30m$

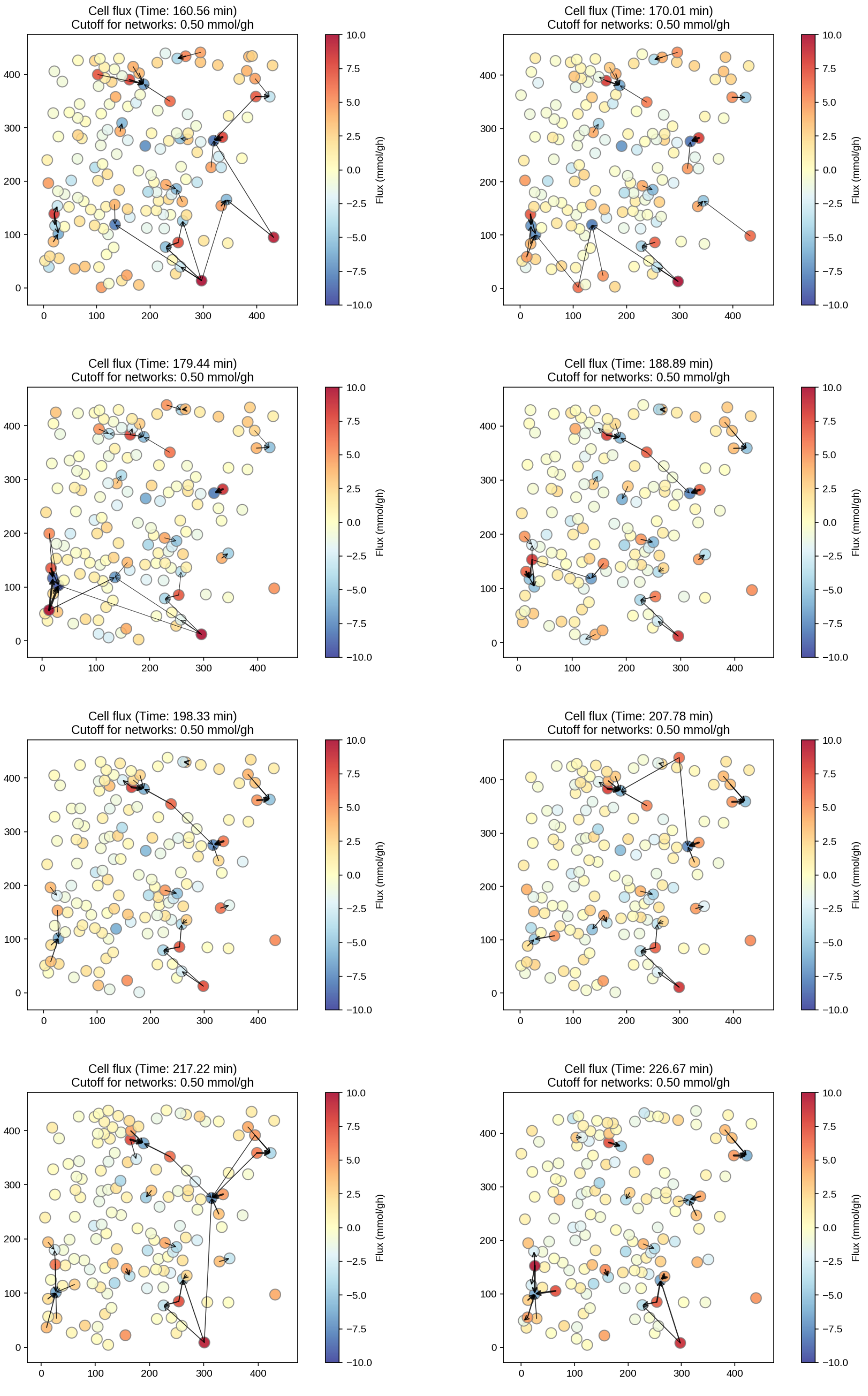

**Figure 3:** Cell fluxes time $2h30m < t < 3h45m$

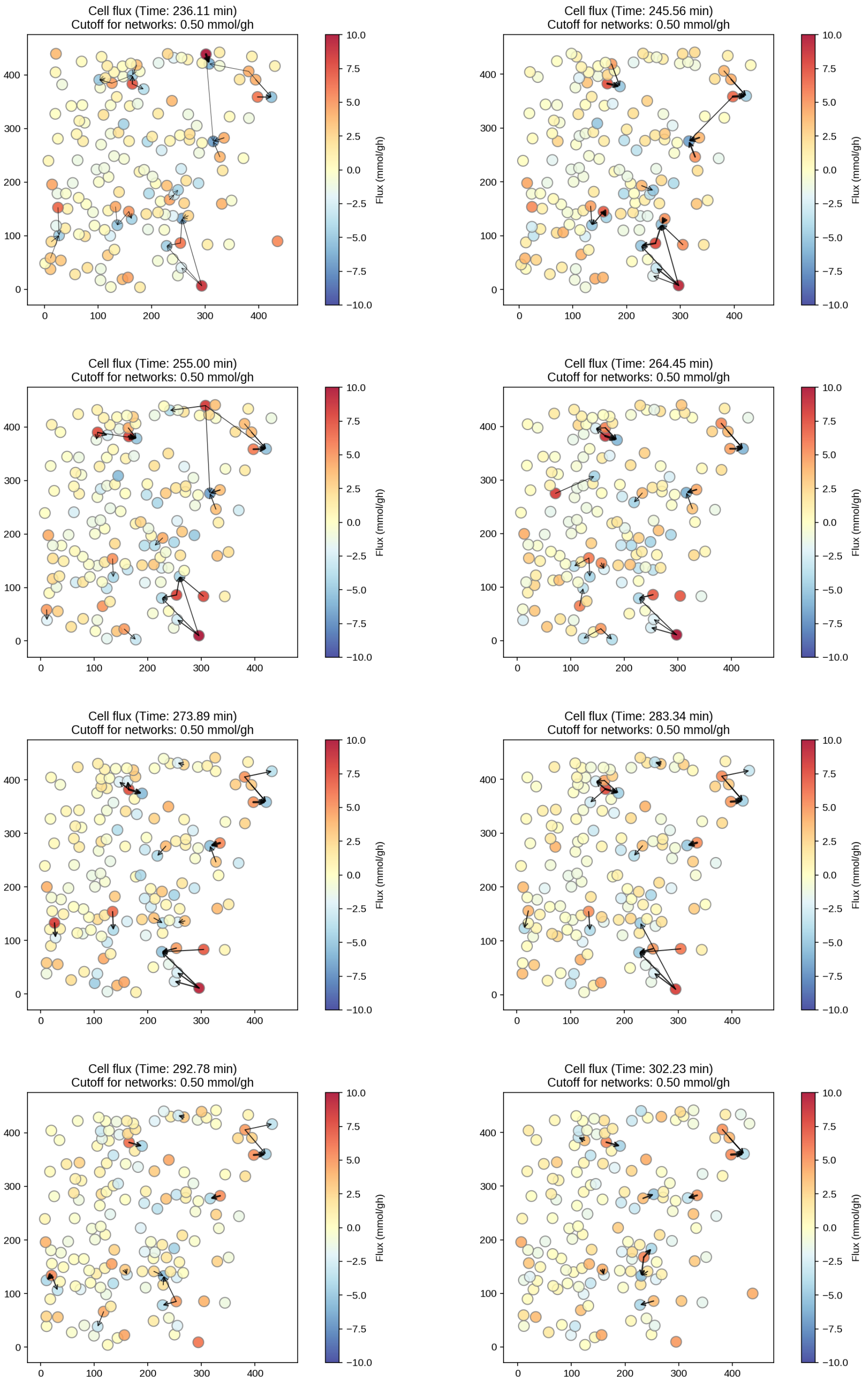

**Figure 4:** Cell fluxes time $3h45m < t < 5h$

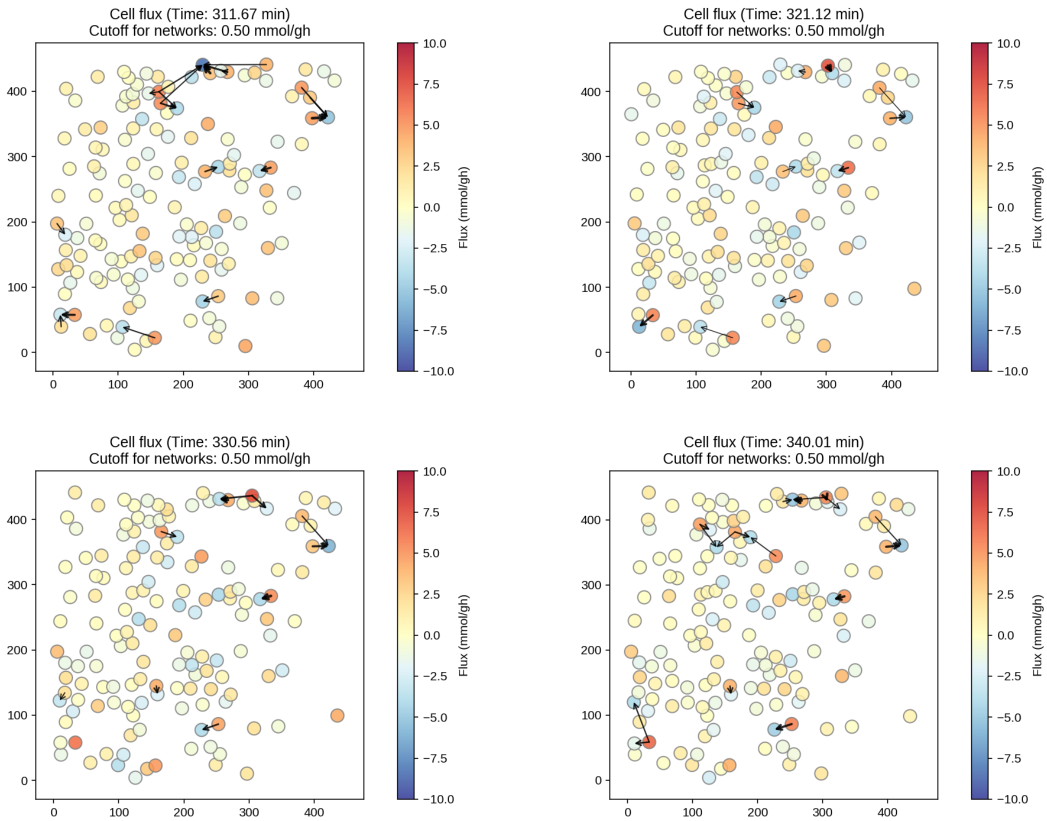

**Figure 5:** Cell fluxes $t > 5\text{h}$

#### 5.2 Errors on recontructed pH at the probes

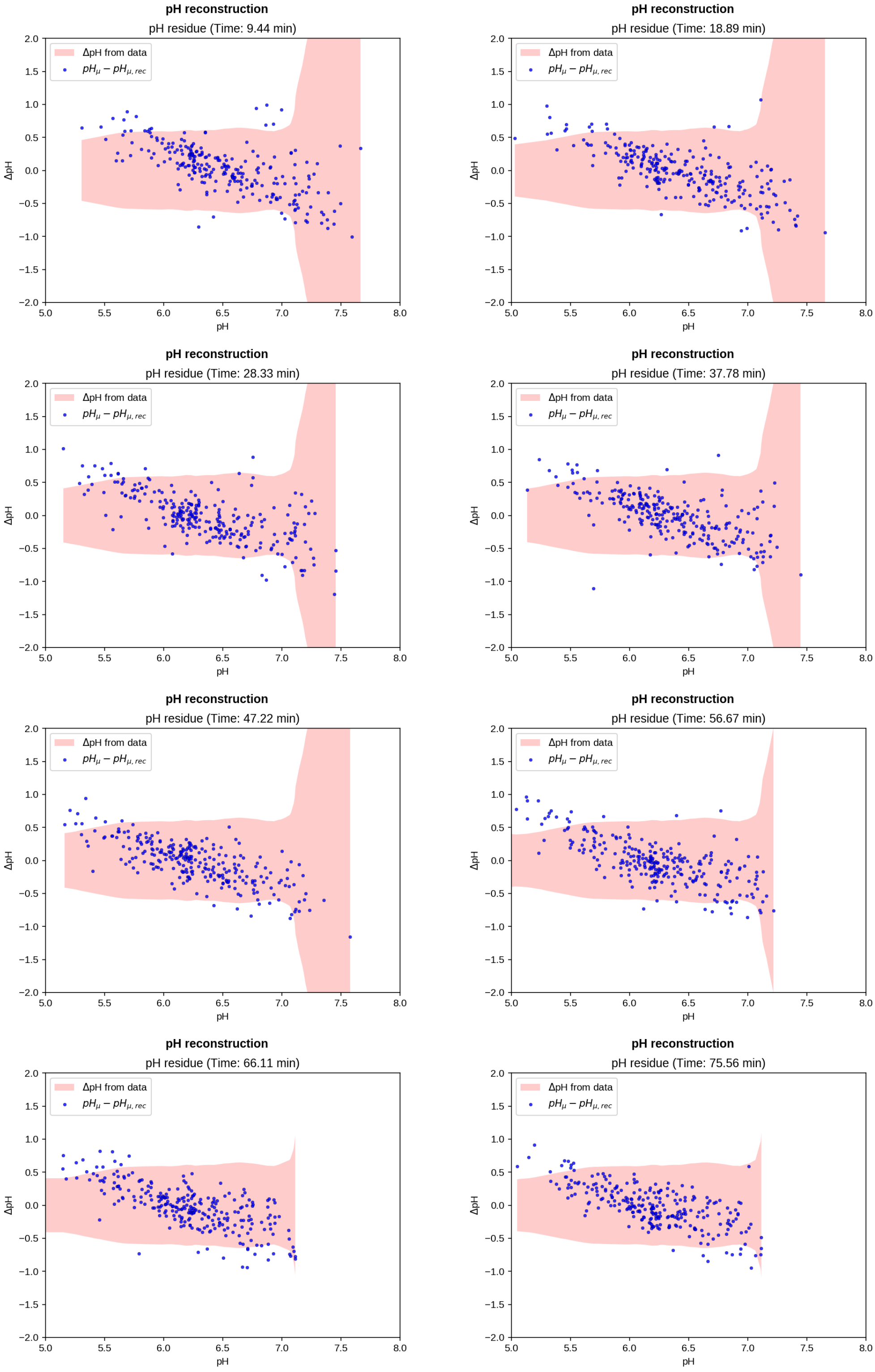

**Figure 6:** Error on reconstructed pH $t < 1h20m$

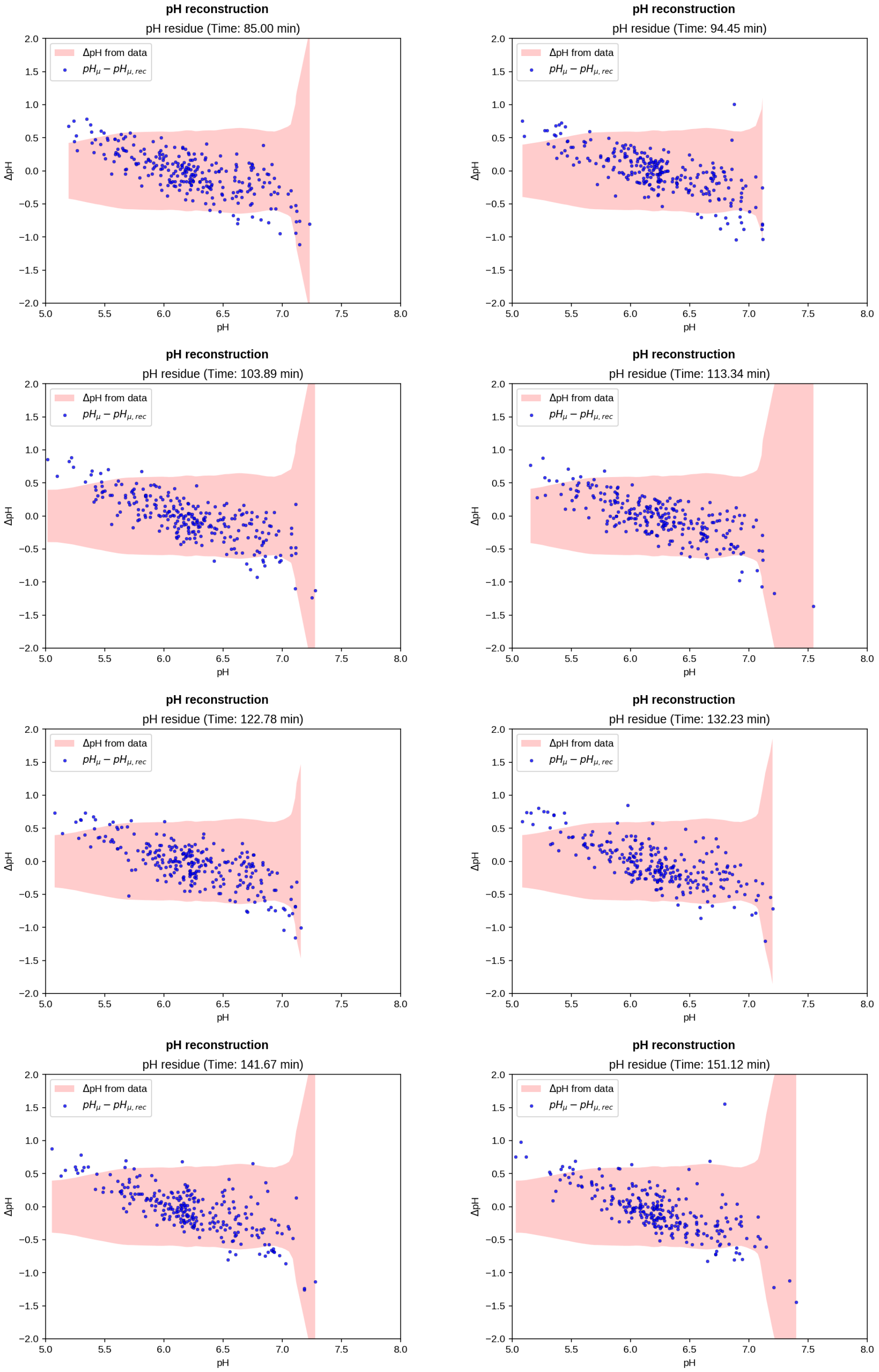

**Figure 7:** Error on reconstructed pH $1h20m < t < 2h30m$

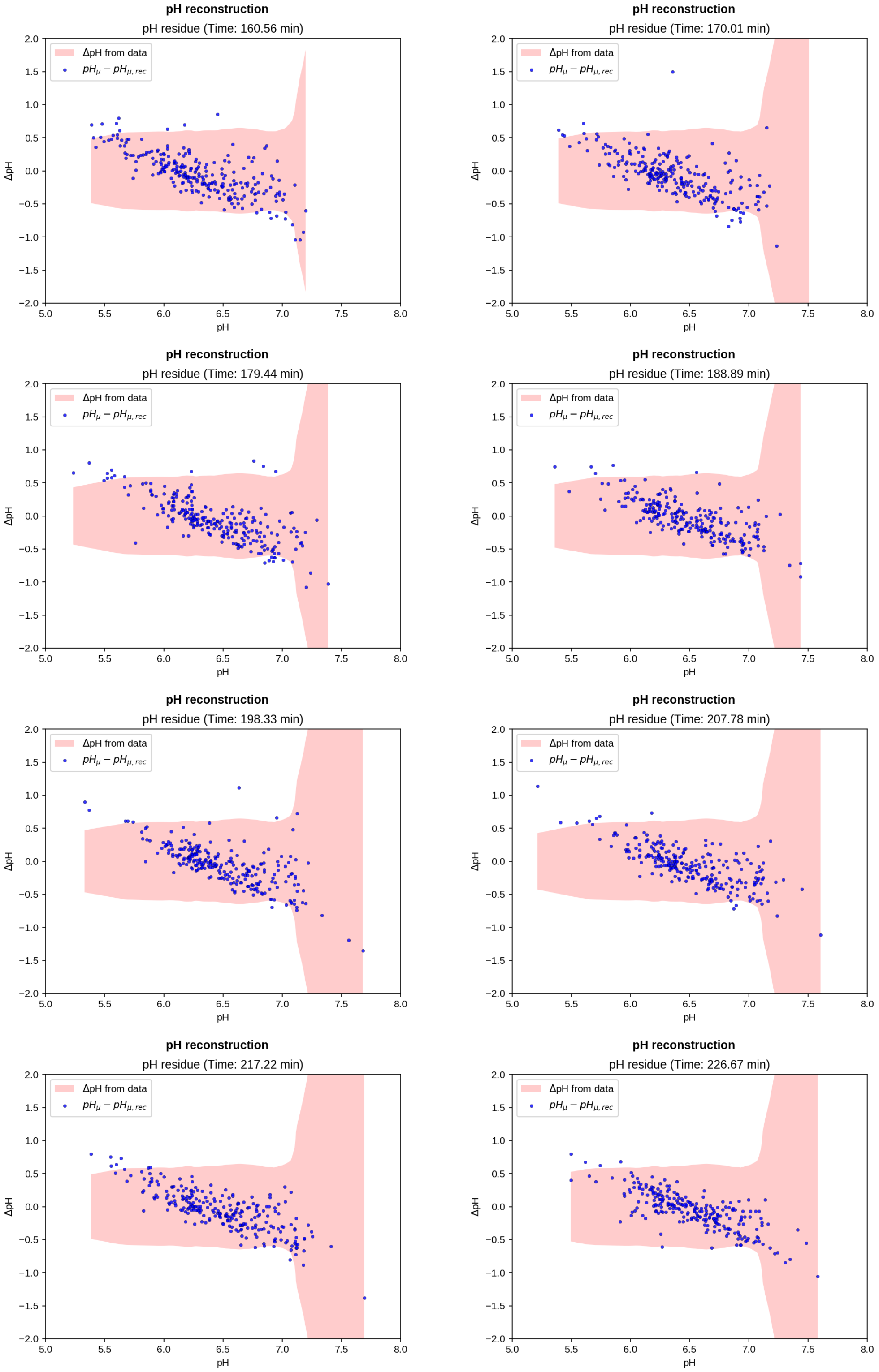

**Figure 8:** Error on reconstructed pH $2h30m < t < 3h45m$

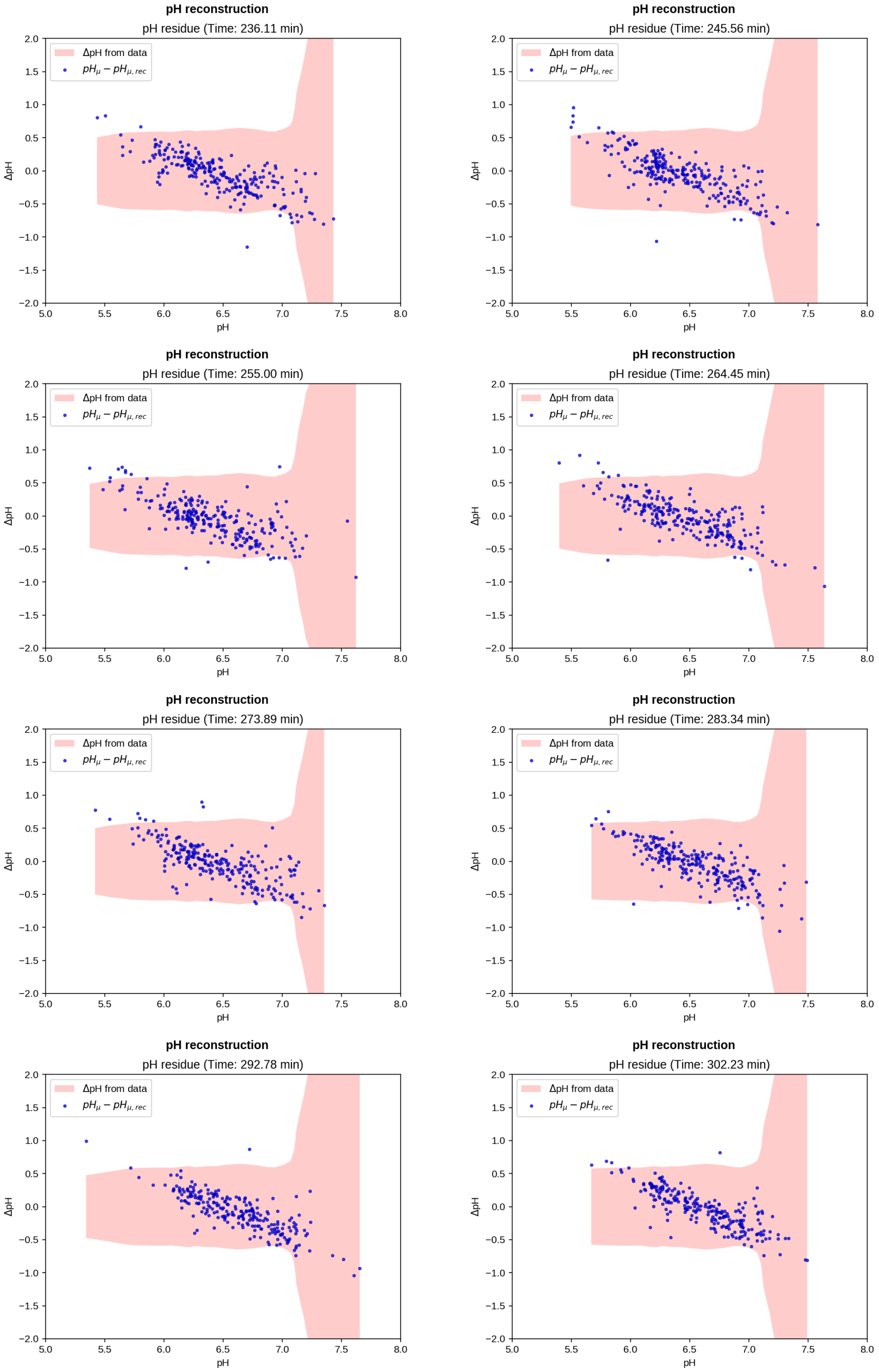

**Figure 9:** Error on reconstructed pH $3h45m < t < 5h$

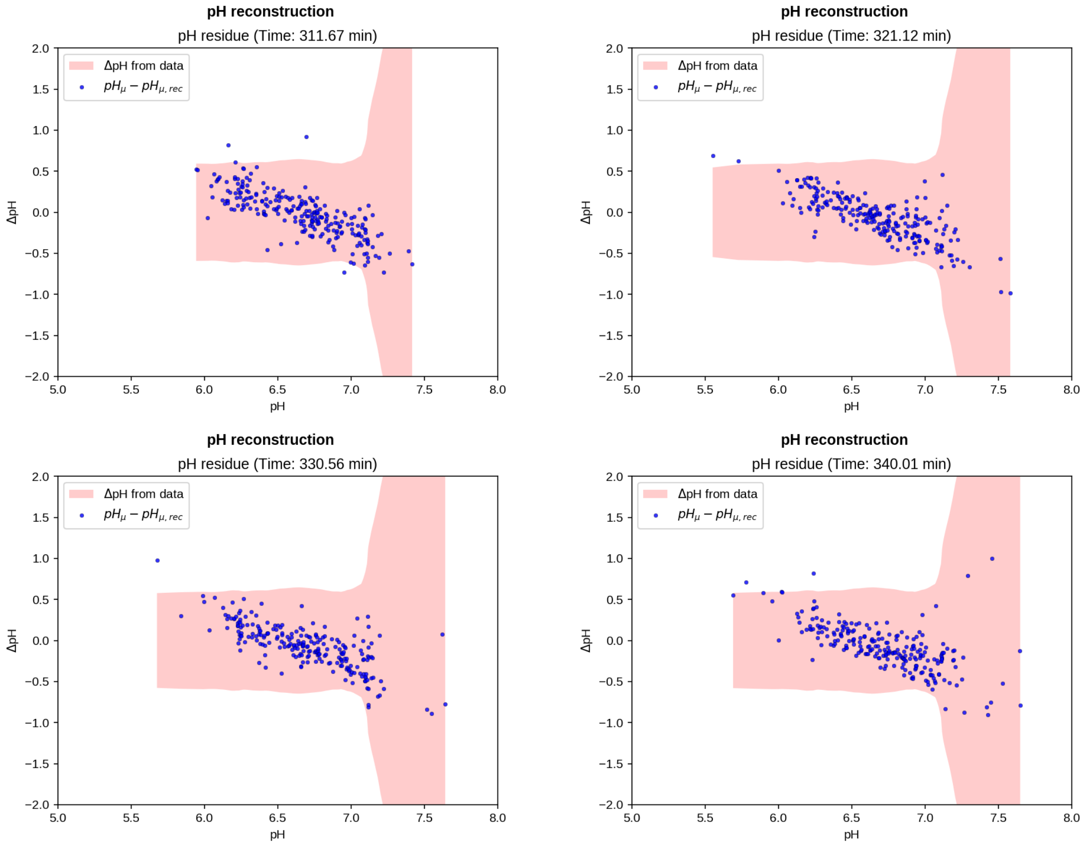

**Figure 10:** Error on reconstructed pH $t > 5h$

### References

[1] Anil Chandra, Saumya Prasad, Francesco Alemanno, Maria De Luca, Riccardo Rizzo, Roberta Romano, Giuseppe Gigli, Cecilia Bucci, Adriano Barra, and Loretta L Del Mercato. Fully automated computational approach for precisely measuring organelle acidification with optical ph sensors. *ACS Applied Materials & Interfaces*, 2022.

[2] Caroline A Schneider, Wayne S Rasband, and Kevin W Eliceiri. Nih image to imagej: 25 years of image analysis. *Nature methods*, 9(7):671–675, 2012.

[3] Frank Y Shih. *Image processing and mathematical morphology: fundamentals and applications*. CRC press, 2017.

[9] David JC MacKay. *Information theory, inference and learning algorithms.* Cambridge university press, 2003.

[10] Frederick James. *Statistical methods in experimental physics.* World Scientific Publishing Company, 2006.
